# Scavenger Cells Failure to Maintain Systemic RNA Homeostasis Causes Epigenetically Inherited Germline Tumors

**DOI:** 10.64898/2026.03.09.710559

**Authors:** Itai Rieger, Yael Mor, Itamar Lev, Anat Nitzan, Charlotte Beata Kong, Sarit Anava, Hila Gingold, Ronen Zaidel-Bar, Oded Rechavi

## Abstract

Temporary disruptions to epigenetic mechanisms can misroute development and permanently alter cell fate. In particular, it was recently shown that transient loss of Polycomb silencing in flies irreversibly reprograms cells toward cancer (Parreno et al. 2024). Whether somatic dysfunction in parents can create multi-generational heritable susceptibility to tumorigenesis is unknown. In eutelic organisms like *Caenorhabditis elegans*, adult somatic cells no longer divide, precluding somatic cancer, yet tumors can still form in the continuously dividing germline. Here, we show that disruption of coelomocytes, somatic scavenger cells, just in *C. elegans* mothers, provokes transgenerationally heritable germline tumorigenesis that persists for multiple generations in genetically wild-type descendants. We found that when the coelomocyte’s phagocytic activity dysfunctions, it impairs clear out of RNA from body fluids, and thus disrupts systemic RNA homeostasis, allowing excess somatic RNAs to access the germline, and leading to widespread transcriptional and small RNA dysregulation and transgenerational loss of germline identity. Converging lines of evidence point towards small RNAs being the heritable agents carrying the pathological information. Together, these findings highlight mechanisms which maintain systemic RNA homeostasis as an important protective barrier against heritable tumorigenesis.

## Introduction

Epigenetic mechanisms enable biological states to persist beyond the conditions that created them (Fitz-James and Cavalli 2022; Ketting and Cochella 2021), raising the possibility that transient physiological events could influence disease risk across generations. However, epigenetic reprogramming and the Weismann barrier (the separation of the soma from the germline) are thought to “reset” parental influences so that the organism can develop according to the hard-wired genetic instruction carried by the germ cells (Heard and Martienssen 2014; Weismann 1893). Whether, nevertheless, epigenetic inheritance contributes to the “Missing heritability”, the portion of inherited trait variation that cannot yet be accounted for by identified DNA differences (Trerotola et al. 2015), remains extremely controversial.

In *Caenorhabditis elegans*, small RNAs mediate potent and heritable changes in gene expression. Numerous studies have shown that small interfering RNAs (siRNAs) and PIWI-interacting RNAs (piRNAs), together with Argonaute proteins and germline-enriched structures such as P granules, can transmit information across multiple generations (Rechavi, Minevich, and Hobert 2011; Ashe et al. 2012; Buckley et al. 2012; Luteijn et al. 2012; Lev et al. 2019; Beltran et al. 2020; Ishikawa and Schumacher 2025). These findings established *C. elegans* as a key model for studying non-DNA-based inheritance.

Disruptions to epigenetic regulation can have profound effects on cell fate. Recent work has shown that transient loss of Polycomb-mediated silencing can irreversibly reprogram cells toward cancer (Parreno et al. 2024), illustrating how brief regulatory failures may generate lasting pathological outcomes. These observations raise an important question: can short-lived physiological disturbances create transgenerationally inherited susceptibility to tumor formation? While parental life experiences, e.g. smoking, can directly influence cancer risk in the developing embryo (Rumrich et al. 2016), transgenerational (in contrast to intergenerational) inheritance of damage for additional generation, unexposed to the parental environment, is unknown.

Systemic homeostasis depends on mechanisms that clear circulating extracellular macromolecules. Coelomocytes are six large scavenger cells that line the pseudocoelom (body cavity) of the *C. elegans* worm. They continuously engulf macromolecules from the pseudocoelomic fluid, performing functions reminiscent of macrophages and liver cells in higher animals (Fares and Greenwald 2001; Schwartz et al. 2010; Zhang, Grant, and Hirsh 2001). Their role in maintaining extracellular homeostasis suggests that they could influence the distribution of circulating RNAs and thereby modulate the regulatory environment encountered by the germline. Supporting this possibility, studies in *Drosophila melanogaster* have shown that hemocytes can process and redistribute double-stranded RNAs (dsRNAs) to promote systemic RNA interference (RNAi), highlighting how scavenger-like cells may actively shape RNA trafficking (Tassetto, Kunitomi, and Andino 2017). However, whether such organism-level regulation affects the progeny is unknown.

Here we describe the discovery that disrupting coelomocyte function is sufficient to generate heritable susceptibility to germline tumorigenesis. We find that loss of coelomocyte-mediated clearance perturbs RNA distribution, destabilizes germline identity, and induces tumor formation that persists across multiple generations in genetically wild-type descendants.

## Results

### Coelomocyte dysfunction compromises germline integrity

To test whether systemic clearance mechanisms influence germline biology, we examined animals lacking *cup-4* (we used two different alleles: *ok837* and *lst1684*), a coelomocyte-specific receptor required for endocytosis (Mergan, Driesschaert, and Temmerman 2023; Patton et al. 2005). We confirmed that *cup-4* is specifically expressed in coelomocytes using single cell data (Ghaddar et al. 2023) (Fig. S1).

Strikingly, *cup-4* mutants exhibited a high incidence of germline tumors (Fig. 1A,B). These structures represent true germline tumors (neoplasms) rather than simple hyperplasia, as they display severely disrupted gonadal architecture and diffuse H2B signal that is no longer confined to discrete nuclei. Tumor frequency increased with age, consistent with progressive germline dysregulation (Garigan et al. 2002; Hughes, Huang, and Kornfeld 2011; Riesen et al. 2014). RNAi-mediated knockdown of *cup-4* similarly elevated tumor incidence, supporting a direct link between coelomocyte function and germline stability (Fig. 1C).

**Fig. 1:**
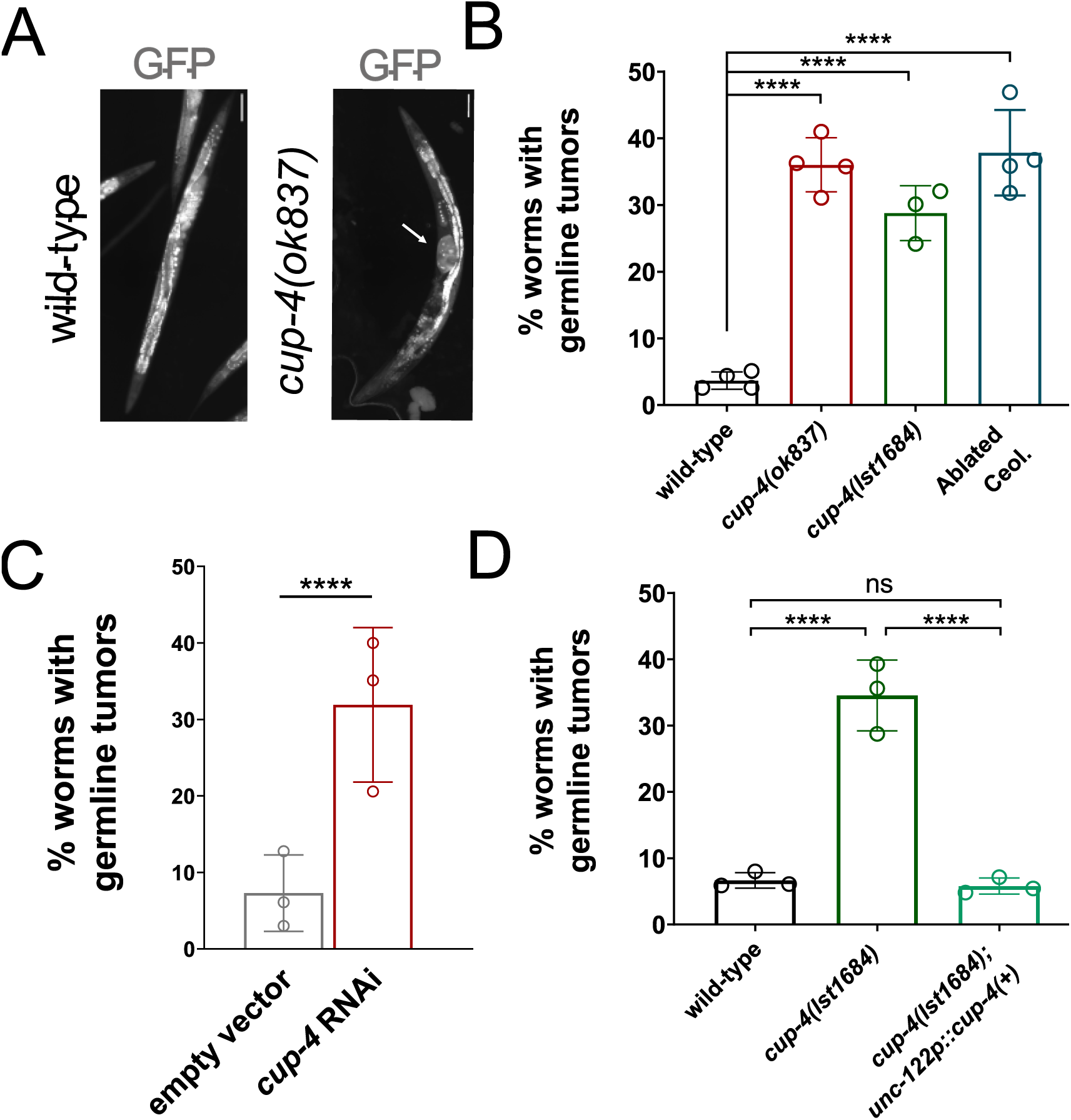
Coelomocyte dysfunction compromises germline integrity. (A) Representative fluorescence images of wild-type and *cup-4* mutants at day 4 of adulthood. White arrow indicates the presence of a germline tumor in *cup-4* animals. Scale bar: 100 µm. (B-D) Quantification of germline tumor penetrance at day 4 of adulthood across indicated genotypes (B+D) or RNAi treatment (C). All strains express the germline marker *mex-5p::GFP*. Data represent means ± SD from 3–4 independent biological replicates. (C) Data is the same as presented in Fig. 2C P0.

Consistent with the established role of coelomocytes in detoxification (Schwartz et al. 2010), we found that *cup-4* mutants were hypersensitive to cadmium exposure (Fig. S2A,B). In addition, these animals were modestly shorter and displayed a Dumpy morphology relative to wild-type controls (Fig. S2C). However, their brood size remained unchanged (Fig. S2D), suggesting that the germline tumors do not arise because of fertility defects. Furthermore, over years of growing *cup-4* mutants, we did not observe a mortal germline phenotype.

To confirm that these phenotypes arise specifically from impaired coelomocyte function, we examined mutants that express a wild-type *cup-4* cDNA under the coelomocyte-specific *unc-122* promoter. Transgenic rescue reduced tumor incidence to wild-type levels and partially restored body length (Fig. 1D, Fig. S2E). Conversely, genetic ablation of coelomocytes, by expressing Diphtheria toxin under the coelomocyte specific *unc-122* gene, phenocopied the mutant animals, resulting in elevated tumor formation (Fig. 1B).

Together, these findings demonstrate that somatic disruption of coelomocyte function is sufficient to compromise germline integrity.

### Somatic coelomocyte dysfunction induces heritable germline tumor formation in genetically wild-type F3 descendants

Because coelomocyte disruption compromised germline integrity, we asked whether these effects were confined to mutant animals or could influence subsequent generations. In *C. elegans* different environmental stresses trigger transgenerational effects (Rechavi, Minevich, and Hobert 2011; Ashe et al. 2012; Buckley et al. 2012; Luteijn et al. 2012; Beltran et al. 2020; Alcazar, Lin, and Fire 2008; Vastenhouw et al. 2006), raising the possibility that somatic physiological disturbances might have lasting heritable consequences.

To test this, we crossed *cup-4* mutants with wild-type animals and monitored genetically wild-type descendants for multiple generations (Fig. 2A). Remarkably, wild-type F3 progeny derived from *cup-4* mutants exhibited a significantly elevated incidence of germline tumors compared to control lineages (Fig. 2B left). Thus, transient disruption of a somatic cell type was sufficient to induce heritable germline tumor formation in otherwise genetically normal descendants.

**Fig. 2:**
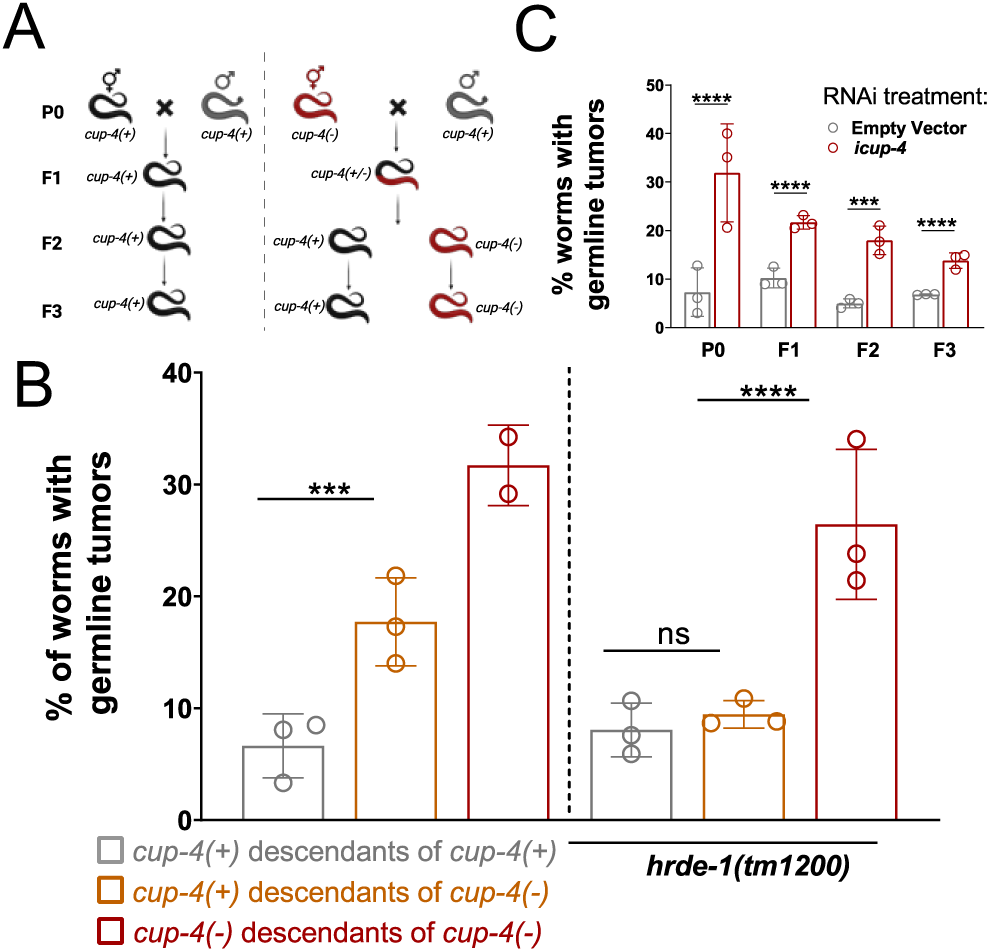
Somatic coelomocyte dysfunction induces heritable germline tumor formation. (A) A schematic diagram of the crosses performed to examine the effects of *cup-4(ok837)* mutation in the ancestry on the phenotype of wild type progeny. On the left: wild type hermaphrodites were crossed to wild type males and their F3 wild type offspring were tested for the presence of germline tumors, for a negative control. On the right: *cup-4(ok837)* mutant hermaphrodites were crossed to wild type males and their F3 wild type (and *cup-4* for positive control) offspring were tested for the presence of germline tumors. (B) Quantification of germline tumor penetrance at day 4 of adulthood across indicated genotypes. All strains express the germline marker *mex-5p::GFP*. Data represent means ± SD from 2-3 independent biological replicates. On the right: worms containing a background mutation for *hrde-1*. (C) Quantification of germline tumor penetrance at day 4 of adulthood across indicated RNAi treatment. All strains express the germline marker *mex-5p::GFP*. Data represent means ± SD from 3 independent biological replicates. Data for P0 is the same as presented in Fig. 1C

We next asked whether this inheritance depended on small RNA pathways known to mediate transgenerational epigenetic effects in *C. elegans*. Strikingly, the increased tumor susceptibility was abolished in an *hrde-1* mutant background (Fig. 2B right), (encoding for the nuclear Argonaute HRDE-1 (Ashe et al. 2012; Buckley et al. 2012; Luteijn et al. 2012)) and thus implicated small RNAs as carriers of the inherited information.

Not all phenotypes associated with coelomocyte dysfunction were transmitted across generations. Hypersensitivity to cadmium was not inherited, and reductions in body length were variable (Fig. S3).

Finally, we observed increased tumor formation in descendants of animals exposed to *cup-4* RNAi (Fig. 2C). Silencing of somatic genes is typically in itself not transgenerationally heritable (Ewe and Rechavi 2023), and therefore these findings, together with the mutant analysis, suggest that transient somatic perturbations can indirectly trigger transgenerational phenotypic consequences that persist beyond the initiating event.

### Coelomocyte dysfunction disrupts germline RNA regulation and cell identity

To understand how somatic coelomocyte dysfunction destabilizes the germline, we looked for changes in the regulatory pathways known to preserve germline identity. Epigenetic mechanisms play a central role in maintaining germ cell fate and preventing inappropriate somatic transcriptional programs (Carpenter et al. 2021; Ciosk, DePalma, and Priess 2006; Knutson et al. 2017; Updike et al. 2014). We therefore profiled small RNAs and mRNAs from dissected gonads of wild-type and *cup-4* mutant animals prior to overt tumor formation. This analysis revealed widespread misregulation of small RNAs in the *cup-4* mutants’ germline, with large numbers of small RNAs both upregulated and downregulated relative to wild type (Fig. 3A). Notably, the downregulated small RNAs were highly enriched for WAGO-1, WAGO-4, CSR-1, and HRDE-1-associated classes (Fig. 3B). In parallel, mRNA sequencing also showed widespread misregulation (Fig. 3C), and uncovered widespread aberrant expression of somatic genes in the *cup-4* germline, a hallmark of germline identity loss and teratoma-like tumor formation (reviewed in (Ul Fatima and Tursun 2020)) (Fig. 3D). These findings align with prior studies linking ectopic somatic gene expression in the germline to failures in epigenetic regulation, including disruption of P granules and small RNA pathways (Carpenter et al. 2021; Ciosk, DePalma, and Priess 2006; Knutson et al. 2017; Tursun et al. 2011; Updike et al. 2014). In particular, Knutson et al (Knutson et al. 2017) sequenced mRNAs from worms lacking P granules that also express somatic genes in the germline. Strikingly, mRNA-seq from dissected gonads of *cup-4* mutants closely recapitulate these findings, with 529 of 607 genes, 87%, overlapping between the sets of upregulated genes. Consistently, enrichment analysis revealed a strong and highly significant overlap between the two datasets (fold enrichment of 1.92, P < 0.0001). Taken together, these data reveal a profound disruption of germline small RNA mediated gene regulation in *cup-4* mutants, accompanied by widespread ectopic somatic gene expression that precedes tumor formation.

**Fig. 3:**
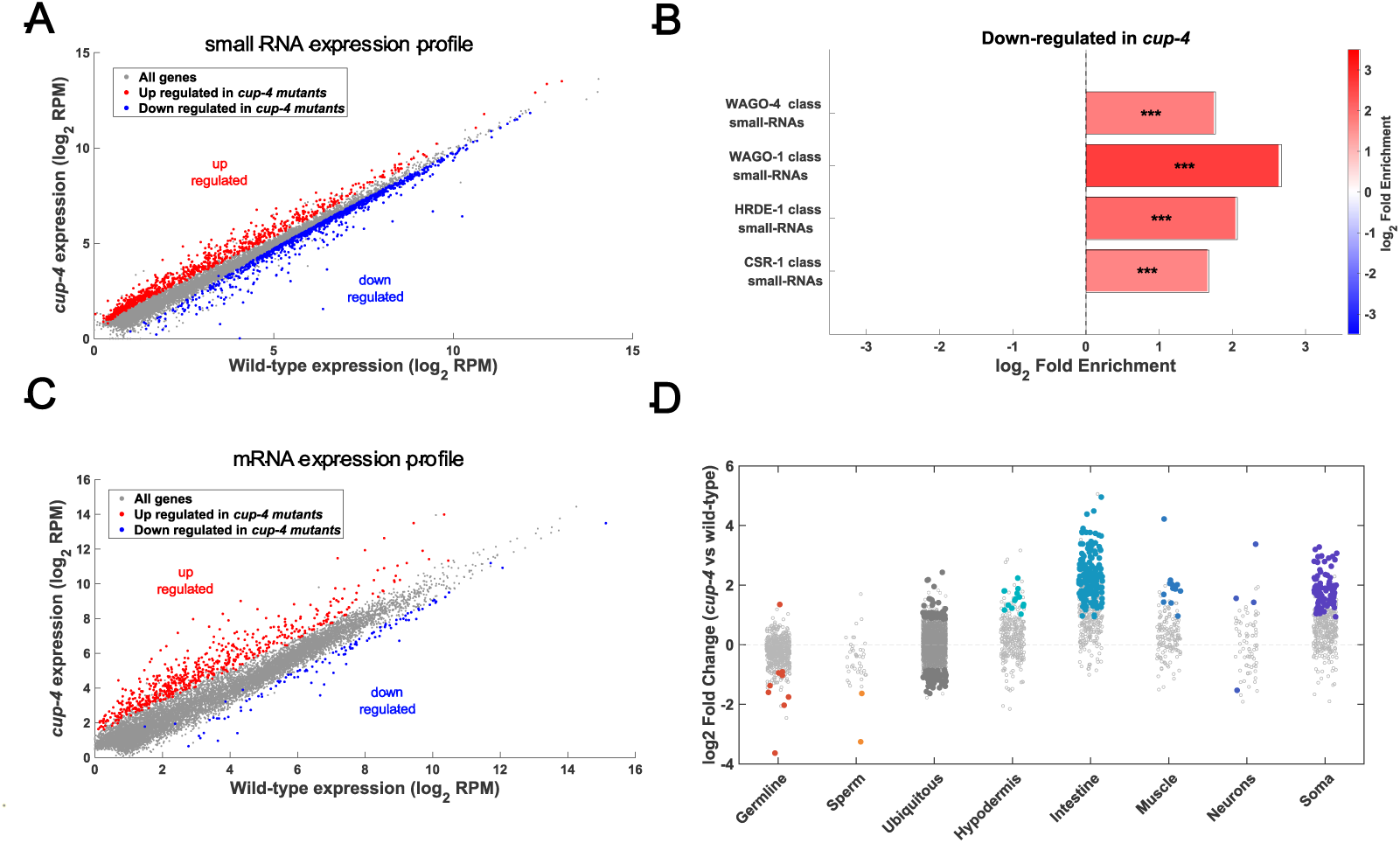
Coelomocyte dysfunction disrupts germline RNA regulation and cell identity. (A+C) Expression of small RNAs (A) or mRNA (C) in gonads of *cup-4(ok837)* mutants (Y-axis) versus wild type worms (X-axis). Shown are averaged expression values (log2 RPM). Each dot represents small RNAs targeting a specific gene (A) or mRNA gene expression (C). Red and blue dots indicate small RNAs that are significantly up- or down-regulated, respectively, in *cup-4* mutants (DESeq2, adjusted p-value < 0.1).(B) X-fold enrichment values (log2, bar graphs and color-coded). Shown are results for genes with significantly downregulated levels in *cup-4(ok837)* mutants’ gonadal small RNA compared to wild type. We tested the enrichment against lists of genes that are known targets of the Argonautes described (Buckley et al. 2012; Claycomb et al. 2009; Gu et al. 2009). P values for enrichment were calculated using 10,000 random gene sets identical in size to the tested group. (D) Log2 fold-change of mRNA expressed in dissected gonads of *cup-4(ok837)* mutants vs. wild-type for genes expressed in different tissues based on tissues specific gene lists (Serizay et al. 2020). Filled colored dots represent significantly differentially expressed genes (DESeq2, adjusted p-value < 0.1).

### Coelomocyte dysfunction alters RNA trafficking between the soma and the germline

Images from previously published data suggested that dsRNA accumulates in the coelomocytes (see figures 6A and 4B from Marré, Traver, and Jose 2016; Wang and Hunter 2017 respectively).We hypothesized that the misregulation of regulatory RNAs in the *cup-4* mutants’ germline (Fig 3A,B), and also the germline tumor formation, result from impaired RNA clearance which leads to dysregulated RNA distribution in the pseudocoelom, and consequently also in the germline. To examine systemic RNA trafficking, we tracked the fate of fluorescently labeled dsRNAs that were injected into either wild-type or *cup-4* animals. In wild-type worms, coelomocytes efficiently internalized dsRNAs, whereas in *cup-4* mutants uptake was reduced. We note that within the same worm, some coelomocytes internalized dsRNAs normally, while others failed completely, for 2 different concentrations of injected dsRNA: 500 and 200 ng/µl (Figure 4A,B). Concomitantly, more dsRNA accumulated in the germline of *cup-4* mutants (Figure 4C). These results raised the possibility that coelomocyte dysfunction disrupts systemic RNA trafficking, increasing RNA accumulation in the germline. If small RNA trafficking is perturbed, one would predict that gene silencing would also be affected. Therefore, we tested RNAi by feeding the worms with dsRNAs targeting both somatic and germline genes (*pos-1* and *Peft-3::mCherry*). In all cases, *cup-4* mutants showed enhanced RNAi phenotype (Eri) (Figure 4D,E), establishing that when RNA clearance by the coelomocytes is disrupted, small RNA dependent gene silencing is altered.

**Fig. 4:**
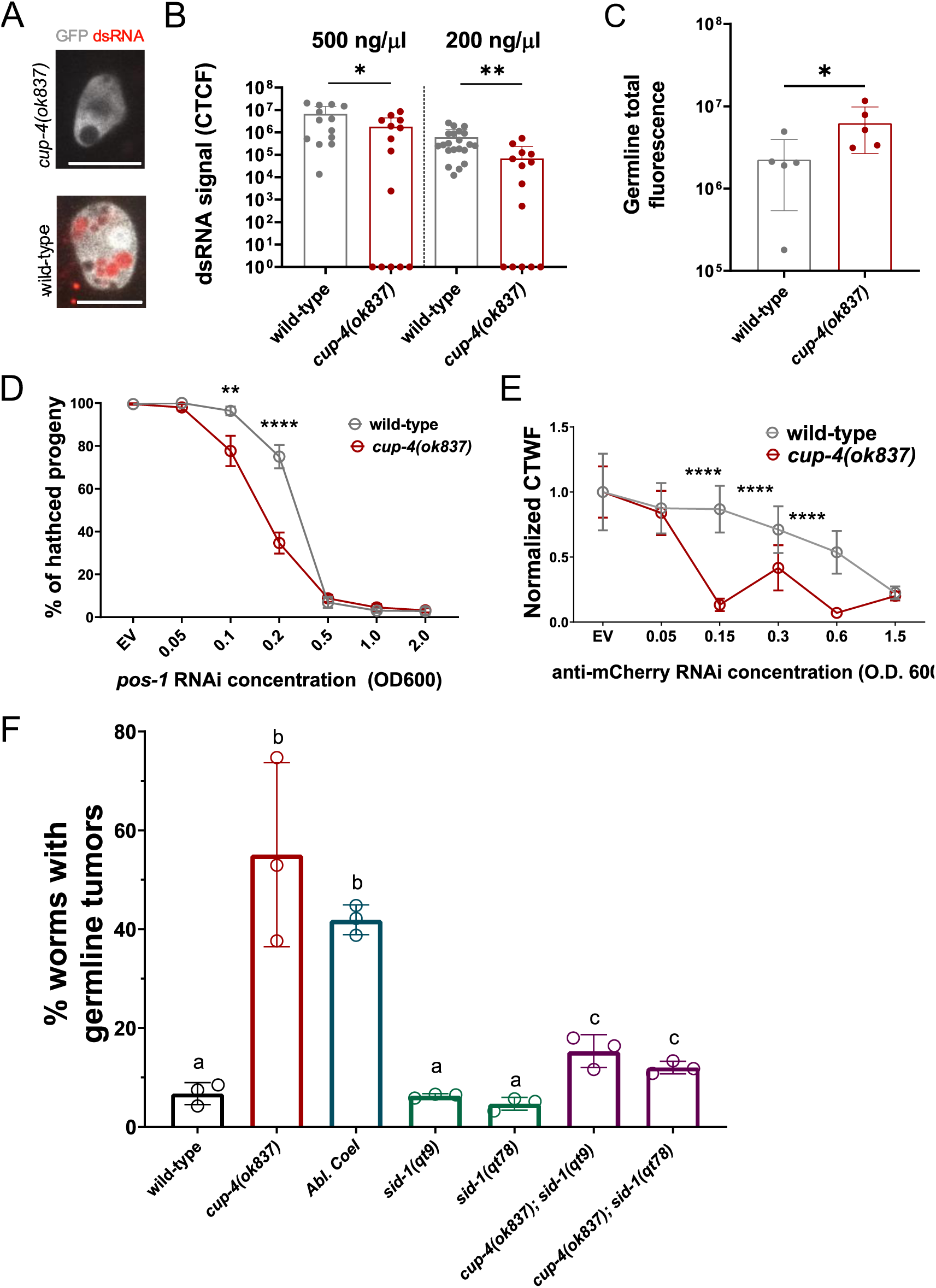
Coelomocyte dysfunction alters RNA trafficking between soma and germline. (A) Representative images of coelomocytes (gray) of *cup-4(ok837)* mutants (top) and wild-type (bottom) hermaphrodites after injection with Cy5-labeled *unc-22* dsRNA (red). Coelomocytes are visualized using the *wyIs629 [Pgcy-8::GCaMP6s; Pgcy-8::mCherry, Punc-122::GFP]* transgene. Scale bar: 10µm (B) coelomocytes corrected total cell fluorescence (CTCF) in wild-type and *cup-4(ok837)* mutants after injection with two different concentrations of tagged dsRNA. (C) total fluorescence in germline of wild-type and *cup-4(ok837)* mutants after injection with 200 ng/µl tagged dsRNA. (D) percentage of hatched progeny in wild-type (gray) and *cup-4(ok837)* (red) mutants after exposure to different concentrations of dsRNA targeting *pos-1* gene. Shown are the mean and SEM of four combined biological replicates. (E) corrected total worm fluorescence (CTWF) of each worm was divided by the mean fluorescence of the control empty vector treated worms. The fluorescence of each worm was obtained automatically using WorMachine (Hakim et al. 2018). Each dot represents mean fluorescence of the indicated genotype; wild type (grey) and *cup-4(ok837)* (red) after exposure to different concentrations of dsRNA targeting *mCherry* gene. All worms express the ubiquitous *eft-3p::mCherry* transgene. Shown are means ±SD of one experiment. Total worm number ranged between 70 and 380 per condition. (F) Quantification of germline tumor penetrance at day 4 of adulthood across indicated genotypes. All strains express the germline marker *mex-5p::GFP*. Data represent means ± SD from 3 independent biological replicates. Conditions labeled with different letters show statistically significant differences in germline tumor percentage (P < 0.0001).

SID-1 (**S**ystemic RNA **I**nterference **D**eficient 1) is a dsRNA transporter, required for intracellular import of circulating dsRNA (Feinberg and Hunter 2003). If, as we hypothesized, increase in the flow of somatic dsRNA to the germline is involved in the non-cell autonomous effects of coelomocytes mutants (e.g. increased germline tumorigenesis), then one would predict that the effects should decrease in *sid-1* mutants. In agreement with this prediction, we found that in *cup-4;sid-1* double mutants (two different *sid-1* alleles were examined: *qt9* and *qt78*), less germline tumors were developed in comparison to *cup-4* single mutants (Fig 4F). Interestingly, disabling *sid-1* did not restore the levels of tumors in *cup-4;sid-1* double mutants’ germline all the way to wild type levels, suggesting that *cup-4* affects germline development in additional, SID-1 independent ways. These results provide a mechanistic basis for how transient somatic dysfunction can expose the germline to regulatory RNAs capable of reshaping heritable outcomes.

## Discussion

In this work, we demonstrate that disruption of a somatic cell type is sufficient to generate heritable susceptibility to germline tumorigenesis. Loss of coelomocyte function disrupts RNA homeostasis by impairing RNA clearance from the pseudocoelom, leading to germline tumors that are heritably transmitted to genetically wild-type descendants through multiple generations in an HRDE-1 and SID-1-dependent manner.

It might be surprising that disruption of just six somatic cells gives rise to heritable tumorigenesis, however, coelomocytes work non-cell autonomously, and accordingly affect many different physiological processes. Similarly, it was previously shown that disruption of individual neurons, which coordinate different bodily functions has heritable consequences, (Szántó et al. 2024; Teichman et al. 2024).

Interestingly, we found that an *indirect* phenotype, the propensity to develop tumors, is epigenetically inherited, whereas susceptibility to cadmium exposure, a phenotype more *directly* linked to coelomocyte function, is not (Fig. 2B, 3SA). In studies of epigenetic inheritance, heritable phenotypes are often sought that closely mirror the original stimulus or treatment, for example when parental stress leads to altered stress responses in the progeny. Our results emphasize that, given the complexity of biological systems, an initial trigger can spark a cascade of events that culminate in phenotypic outcomes that are not obviously or directly related to the original perturbation. This consideration may therefore be important when interpreting and designing studies of heritable epigenetic effects.

Moreover, we acknowledge the possibility that disrupting coelomocytes functions misregulates germline processes (leading to heritable tumorigenesis) in different ways, not exclusively because of disrupted RNA homeostasis. For example, inappropriate pseudocoelom clearance might generate a stress response, and different types of stress were shown to cause transgenerational effects (e.g. *39*) (although increased heritable tumorigenesis has not been described to our knowledge). We did not observe any clear signature of stress in the germline transcriptome of *cup-4* animals, for example, none of the following stress-related genes: *daf-16, hsf-1, skn-1, nhr-49, hlh-30, daf-2, daf-7,* and *daf-12* were misregulated in the *cup-4* gonadal transcriptome. However, one cannot rule out the possibility that some stress pathway activation went undetected. We note that adding a *sid-1* mutation in the background of *cup-4* mutated animals did not completely restore tumor levels to wild-type level (Fig. 4E), suggesting indeed that additional signals could contribute to the process.

Eutelic organisms like *C. elegans*, which have a fixed number of somatic cells that cease to divide in the adult, are cancer free, since somatic perturbations cannot propagate by clonal expansion. Nevertheless, the worms can develop tumors in their germ cells, which continue to proliferate. Although such germline tumors in *C. elegans* are not malignant, they expose core mechanisms by which proliferating cells actively suppress inappropriate differentiation programs, and loss of cell identity is a defining feature of many cancers. Many pathways that prevent germline tumor formation, including small RNA mediated silencing (investigated here), translational repression (Ciosk, DePalma, and Priess 2006), and chromatin-based mechanisms (Patel et al. 2012; Tursun et al. 2011), are frequently disrupted in human cancers (Darwiche 2020; Flavahan, Gaskell, and Bernstein 2017; Scaffidi and Misteli 2010). Notably, genes upregulated in *cup-4* mutants are significantly enriched for *C. elegans* orthologs of human cancer driver genes (Cerón 2023) (fold enrichment of 3.09, P < 0.0001), suggesting activation of a conserved tumor-associated transcriptional program. Future studies are required to understand if in humans RNA disruption in parents can lead to increased tumorigenesis in the descendants.

## Methods

### Worm cultivation and strains

All worms were grown at 20° C on Nematode Growth Medium (NGM) plates seeded with OP50 bacteria. Worm strains: N2, SX1263: *mjIs134 [mex-5p::gfp::h2b::tbb-2 II]*, RB950: *cup-4(ok837) III*, BFF67: *mjIs134 [mex-5p::gfp::h2b::tbb-2 II]; cup-4(ok837) III*, LSC1961: *cup-4(lst1684) III*, LSC1963: *cup-4(lst1684) III; lstEx1065[unc-122p::cup-4::cup-4 3’UTR + myo-2p::mCherry]*, BFF426: *mjIs134 [mex-5p::gfp::h2b::tbb-2] II; cup-4(lst1684) III*, BFF493: *mjIs134 [mex-5p::gfp::h2b::tbb-2] II; cup-4(lst1684) III; lstEx1065[unc-122p::cup-4::cup-4 3’UTR + myo-2p::mCherry],* BFF548: *mjIs134 [mex-5p::gfp::h2b::tbb-2] II; cdIs32 [unc-122p::DT-A(E148D) + myo-2p::GFP + unc-119(+)]*, BFF48: *mjIs134 [mex-5p::gfp::h2b::tbb-2] II; hrde-1(tm1200)* III, *BFF432: mjIs134 [mex-5p::gfp::h2b::tbb-2] II; hrde-1(tm1200) III cup-4(ok837) III*, BFF56: *SX1263[mjIS134 II[mex-5::gfp::h2b::tbb-2]] II; sid-1(qt9) V*, BFF427: *mjIs134 [mex-5p::gfp::h2b::tbb-2] II; sid-1(qt78) V*, BFF494: *mjIs134 [mex-5p::gfp::h2b::tbb-2] II; cup-4(lst1684) III; sid-1(qt9) V*, BFF433: *mjIs134 [mex-5p::gfp::h2b::tbb-2] II; cup-4(lst1684) III; sid-1(qt78) V*, DCR3055: *wyIs629 [Pgcy-8::GCaMP6s; Pgcy-8::mCherry, Punc-122::GFP]*, BFF86: *wyIs629 [Pgcy-8::GCaMP6s; Pgcy-8::mCherry, Punc-122::GFP];cup-4(ok837) III*, EG7841: *oxTi302 [eft-3p::mCherry::tbb-2 3’UTR + Cbr-unc-119(+)]*, BFF404: *oxTi302 [eft-3p::mCherry::tbb-2 3’UTR + Cbr-unc-119(+)]; cup-4(ok837) III*

### Tagged dsRNA

Extracted RNA was reverse transcribed using SuperScript™ III Reverse Transcriptase to create cDNA. The *unc-22* with flanking T7 promoters was amplified using Phusion polymerase, NEB, and then cloned into pGEM-T Vector. Then, using TranscriptionAid T7 High Yield Transformation kit, labeled UTPs (Cyanine 5-Aminoallyluridine-5’-Triphosphate – Trilink cat# N-5108) were incorporated to transcribe the final tagged dsRNA. Two samples with different RNA concentrations were created – 200 and 500 ng/µl.

### Injection of tagged dsRNA

Adult worms (1 day after the L4 larval stage) carrying a GFP reporter in their coelomocytes (transgene: *wyIs629 [Pgcy-8::GCaMP6s; Pgcy-8::mCherry, Punc-122::GFP]*) were injected with 500 or 200 ng/µl of tagged dsRNA into the body cavity past the bend of the posterior arm of the gonad (near the worm’s tail). Worms were then imaged using Nikon Ti-2 eclipse microscope equipped with a 100X CFI Plan-Apo 1.45 NA objective (Nikon, Tokyo, Japan) and CSU-W1 spinning-disk confocal head (Yokogawa Corporation,Tokyo, Japan).

### CdCl2 plates

CdCl2 was added to NGM at a final concentration of 50μM.

### Gonads dissection for RNA sequencing

wild-type N2 and *cup-4* Hermaphrodites were collected on the first day of adulthood, washed 4 times with M9 buffer, and mounted on a microscope agarose slide with egg buffer (1mM HEPES, 5mM NaCl, 1mM MgCl2, 1mM CaCl2, 1mM KCl and 20% tween-20) containing 2mM levamisole. Worms were dissected right under the pharynx or on the tail with a gauge needle, and the emitted gonads separated from the body. Gonads were transferred from the slide into an Eppendorf tube on ice prior to the addition of 300μl Trizol (Life Technologies).

### RNA extraction

3 freeze/thaw cycles - −80 °C for 30 minutes followed by 15 minutes vortex at room temperature. 1 volume of chloroform was added to 5 volumes of Trizol+gonads. The mix was put in a pre-spun 2mL Heavy Phase-Lock tube and centrifuged at 16000 g for 5 minutes at 4 °C. The aqueous phase was transferred to new pre-spun 2mL Heavy Phase-Lock tube, and 1 volume of Phenol:Chloroform:Isoamyl alcohol (sigma) was added per 1 volume of aqueous phase. The tube was centrifuged at 16000 g at room temperature. Aqueous phase was transferred to Eppendorf tube and precipitated with 1 volume of isopropanol and 1.3 µL glycogen (20 µg/µL). the tubes were put for 30 minutes at −20 °C before centrifugation at 16000 g at 4°C. Pellet was washed with 900µL of cold 70% EtOH, and left at room temperature for 20 minutes before being left at −20°C over night. The next morning, the tube was centrifuged at 16000 g for 10 minutes, and the pellet was washed again with cold 70% EtOH. Tubes were centrifuged at 16000 g for 10 minutes, all EtOH was removed and pellet was resuspended in 12 µL of warm (70°C) ddH2O. RNA concentration was determined using Qbit and RNA quality was tested using Agilent 2200 TapeStation.

To ensure the capture of small RNA regardless of their 5’ phosphorylation status, 150-1000 ng from each sample was treated with RNA 5’ polyphosphatase. Concentrations and quality were assessed using Qubit and Agilent 2200 TapeStation respectively.

### small RNA libraries

The NEBNext® Multiplex Small RNA Library Prep Set for Illumina® from New England Biolabs® was used for small RNA library preparation, following the manufacturer’s protocol. The concentration of the samples was determined using Qubit, and their quality was assessed using an Agilent 2200 TapeStation. Subsequently, the samples were pooled and electrophoresed on a 4% agarose E-Gel from Life Technologies. Bands ranging in length from 140 to 160 nt were carefully excised and purified using the MiniElute Gel Extraction Kit (QIAGEN). The purified samples were once again assessed for quality and concentration using the Agilent 2200 TapeStation. Libraries were sequenced on an Illumina NextSeq500 sequencer.

### mRNA libraries

The NEBNext® Ultra II Directional RNA Library Prep Kit for Illumina® from New England Biolabs® was used for mRNA library preparation, following the manufacture’s protocol. The concentration of the samples was determined using Qubit, and their quality was assessed using an Agilent 2200 TapeStation. Samples were then pooled together prior to sequencing on an Illumina NextSeq500 sequencer.

### small RNA seq analysis

The Illumina *.fastq output files were first assessed for quality, using FastQC(Andrews 2010). The files were then assigned to adapters clipping using Cutadapt (Martin 2011). Next, the clipped reads were aligned against the *ce11* version of the C. elegans genome using ShortStack (Shahid and Axtell 2014). We counted reads which align in antisense orientation to genes, using the python-based script HTSeq-count (Anders, Pyl, and Huber 2015) and the corresponding Ensembl-provided gff file (release-95). We then assigned the summarized counts for differential expression analysis using the R package DESeq2 (Love, Huber, and Anders 2014) and limited the hits for genes which were shown to have an FDR < 0.1.

### mRNA seq analysis

mRNA libraries were first assessed for quality using the FastQC tool (Andrews 2010) and were then aligned to *ce11* version of the genome using HISAT2 (Kim, Langmead, and Salzberg 2015). The aligned reads were then counted using the python-based script HTSeq-count (Anders, Pyl, and Huber 2015) and the Ensembl-provided gff file (release-95). Next, the samples were compared for differential expression using the R package DESeq2 (Love, Huber, and Anders 2014). Genes were regarded as differentially expressed if they pass the criterion of FDR < 0.1.

### germline tumor assessment

Worms (all expressing *mjIs134 [mex-5p::gfp::h2b::tbb-2] II)* were age synchronized by limiting 2 hours of egg laying. When reached adulthood, 60 worms per plate (two plates per condition) were transferred every other day to a new plate to avoid starvation. When reached day 4 of adulthood about 120 worms per condition were immobilized with 5 mM levamisole and mounted on 2% agarose pads and fluorescence Images were taken using either Olympus IX83 motorized inverted wide-field microscope or Olympus BX63 motorized upright wide-field microscope at 10x magnification. Tumor presence was analyzed using Fiji/ImageJ software and conditions were blinded from the investigator using the software DoubleBlind. Tumors were scored based on clear disruption of normal gonadal architecture, characterized by disorganized cellular overgrowth. Using a germline nuclear marker, normal germlines exhibit discrete nuclear-localized signal, whereas tumors display multiple densely packed nuclei accompanied by diffuse, hazy H2B signal that is no longer confined to distinct nuclei, indicating loss of normal tissue organization. Tumor incidence was compared using Cochran–Mantel–Haenszel tests stratified by experimental repeat. P-values were adjusted for multiple comparisons using the Holm method.

### Length assessment

worms were aged synchronized by limiting 2 hours of egg laying. When worms reached day1 of adulthood they were washed and immobilized using 5mM sodium azide onto agarose plates per Wormachine protocol (Hakim et al. 2018). Plates were imaged using either Olympus IX83 motorized inverted wide-field microscope or Olympus BX63 motorized upright wide-field microscope at 4x magnification. Length analysis was performed by Wormachine.

### Transgenerational crossed assays

wild-type (for germline tumors: SX1263 – germline fluorescent reporter) males were crossed to L4 wild-type hermaphrodites (control) and *cup-4* (BFF67 for germline tumor experiments) hermaphrodites (P0 generation of the cross, see scheme in Fig. 2A). Individual hermaphrodites were transferred to new plates 24 hours after the cross, and checked for males in the population 3 days later, to establish that they were successfully crossed. L4 F1 progeny of successfully crossed P0 were transferred to new plates. 5 days later F2 worms were put on individual plates (40 from the cross with the *cup-4* genotype and 10 for the control) for 4 hours. After 4 hours worms were taken for *cup-4* genotyping. F3 descendants were then distributed to new plates based on their mother’s genotype (overall 3 conditions: wild-type control from the control cross, wild-type descendants of *cup-4* from the *cup-4* cross and *cup-4* mutants from the *cup-4* cross). F3 worms were subsequently tested for either germline tumors or brood size on cadmium plates or length. This procedure was also done for worms with *hrde-1(tm1200)* mutation in the background.

### Brood size experiment

L1 hermaphrodites were put individually on a plate that either contained or did not contain cadmium. Hermaphrodites were transferred to new OP50 seeded plates since a day after the L4 stage and every 24 hours for a total of 8-10 days. The progeny was quantified on each plate 3 days after the hermaphrodite was there. Hermaphrodites who died with 4 days or less of egg laying were discarded from analysis.

### Lifespan experiment

30 L1 hermaphrodites were put on a plate that either contained or did not contain cadmium. Living hermaphrodites were transferred to new OP50 seeded plates since a day after the L4 stage and every 24 hours. A worm was considered dead if it did not move or react to a touch by a worm pick. After day 10 of adulthood worms were checked for viability everyday but were transferred to new plates every other day.

### RNAi Treatment

HT115 bacteria that transcribe dsRNA targeting *mCherry*, *pos-1*or *cup-4*, were grown in Carbenicillin-containing LB (100 μg/ml) and were then seeded on NGM plates that contain Carbenicillin (25 μg/ml) and IPTG (1 mM). The plates were seeded with bacteria 24 hours prior to their use. All silencing quantification were done either by code or after the person quantifying was blinded to which images/worms are from which condition.

### ipos-1 assay

worms were put on different concentrations of RNAi bacteria at the L4 larval stage. *ipos*-*1* bacteria were diluted in empty-vector to create different concentrations so that the total bacteria was the same in all plates and the worms experienced the same dietary conditions. 24 hours later the worms were removed from the plate, and 24 hours after that, the numbers of hatched progeny and the number of unhatched eggs were counted for each plate.

### imCherry assay

worms were put on different concentrations of RNAi bacteria at the L4 larval stage. *imCherry* bacteria was diluted in empty-vector to create different concentrations so that the total bacteria was the same in all plates and the worms experienced the same dietary conditions. Worms were left for 24 hours to lay eggs. Next generation was put on new RNAi plates on day 1 of adulthood for 2 hours to lay eggs. Next generation was imaged as young adults on 2% agarose plates. Quantification for fluorescence was done by WorMachine (Hakim et al. 2018).

### icup-4 assay

P0 worms were synchronized on RNAi bacteria for 2 hours. Progeny were either kept on RNAi until imaged for germline tumors on day 4 of adulthood or washed and bleached on day 2 of adulthood to generate RNAi clean F1 generation. F1 generation was either imaged on day 4 of adulthood for germline tumors or put on separate plates for 2 hours on day 2 of adulthood to generate the F2 generation. F2 generation followed the same treatment to be imaged and generate F3. F3 were imaged on day 4 of adulthood for germline tumors.

### Statistics

ns: P > 0.05, *: P ≤ 0.05, **: P ≤ 0.01, ***: P ≤ 0.001, ****: P ≤ 0.0001. Tumor incidence was compared between treatment and control animals using Cochran–Mantel–Haenszel tests stratified by experimental repeat, P-values were adjusted for multiple comparisons using the Holm method. RNAi silencing for both *pos-1* and *mCherry,* length and brood size were compared using Two Way ANOVA test with Sidak’s post-hoc correction for multiple comparisons. Total dsRNA fluorescence within the coelomocytes was compared between genotypes using an unpaired, two-tailed t-test. The efficiency of dsRNA accumulation within the germline was compared between wild-type and *cup-4* mutants using a Mann-Whitney U test. Life span was compared using Gehan-Breslow-Wilcoxon test.

## Acknowledgments

We thank all the Rechavi lab members for their helpful comments and fruitful discussions. We thank Dr. Liesbet Temmerman for providing strains containing *cup-4(lst1684)*. Some strains were provided by the CGC, which is funded by NIH Office of Research Infrastructure Programs (P40 OD010440). I.R. is supported partly by a fellowship from the Prajs-Drimmer Institute. O.R. is grateful for funding from the European Research Council (no. 819151), the Eric and Wendy Schmidt Fund for Strategic Innovation (Schmidt Sciences Polymath Award no. 0140001000), the Kahn Foundation (grant no. 0604918421) and the Deutsche Forschungsgemeinschaft (grant no. 0604918111). Pre-submission review was conducted using qed Science (https://www.qedscience.com). Data availability: Raw and processed sequencing files are available under GEO accession number GSE325231. Competing interest: the authors declare no competing interests.

---

**Fig. S1:**
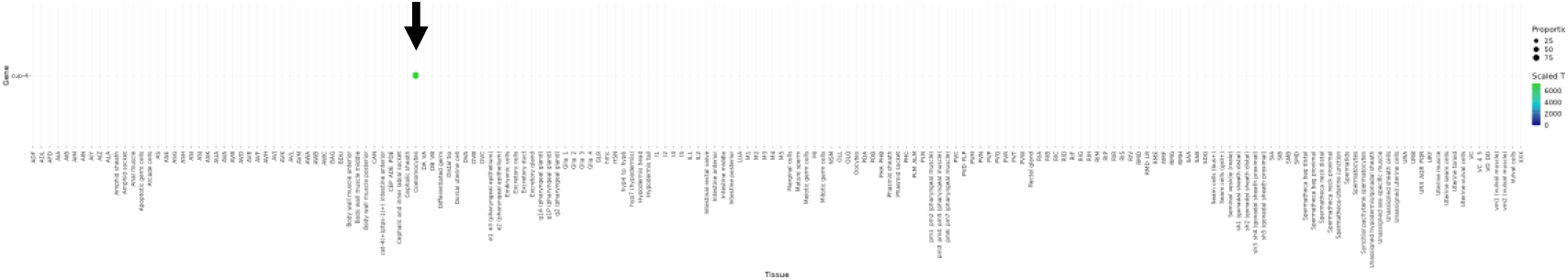
*cup-4* expression across *C. elegans* cell types Data based on work done by Ghaddar et al. (Ghaddar et al. 2023). Black arrow points at the coelomocytes.

**Fig. S2:**
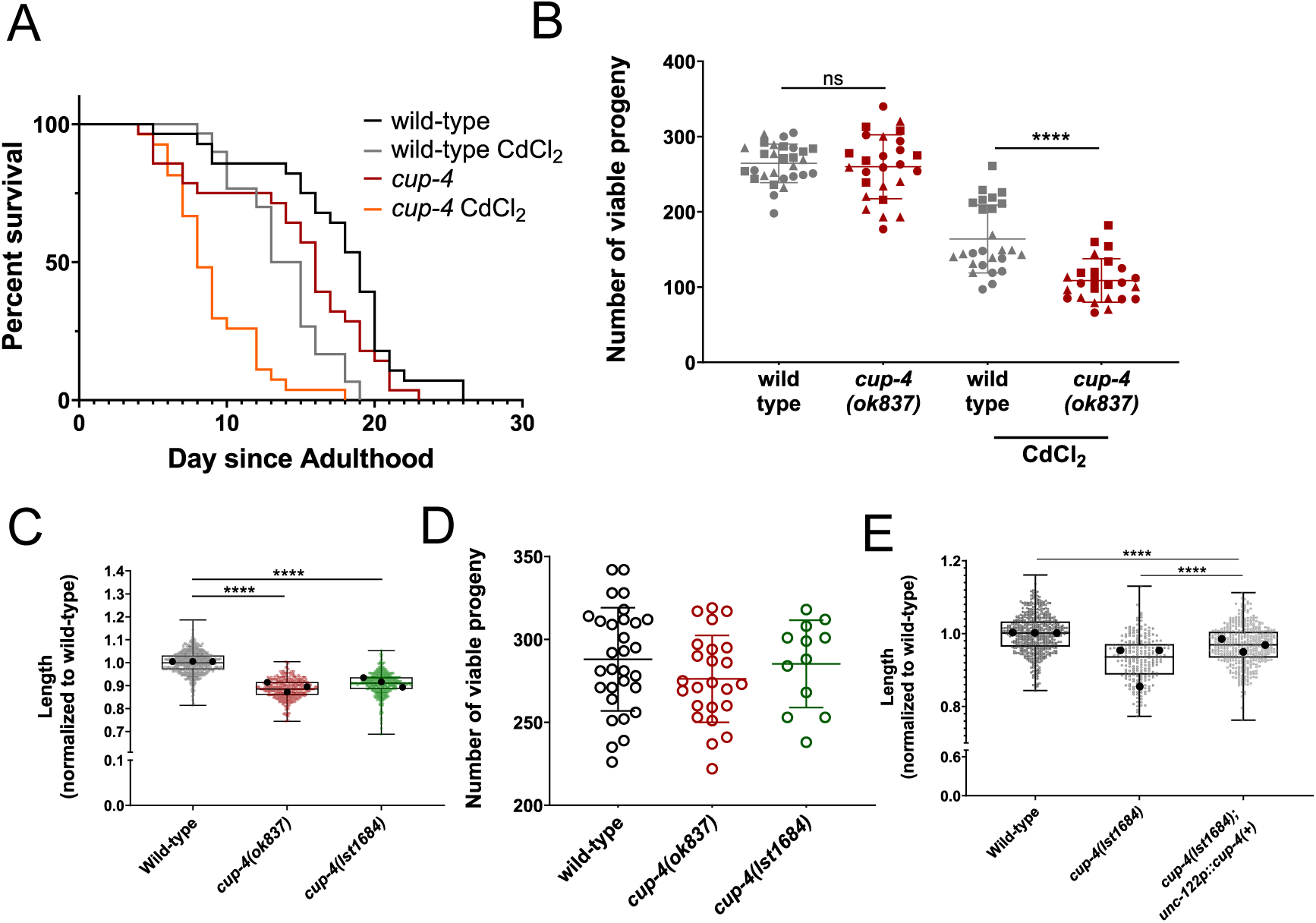
*cup-4* mutants are more sensitive to cadmium exposure and are shorter in length (A) Survival analysis of wild-type and *cup-4(ok837)* mutant strains (n = 30 per group) cultured in the presence or absence of 50 µM CdCl₂. Wild-type animals lived significantly longer than *cup-4* mutants under cadmium stress (P < 0.0001, Gehan-Breslow-Wilcoxon test). (B) Total progeny per individual hermaphrodite (wild-type vs. *cup-4(ok837)*) cultured in the absence (left) or presence (right) of 50 µM CdCl₂. Individual data points represent single hermaphrodites’ brood size, with shapes indicating three independent experimental replicates. (C) Relative body length of wild-type and two *cup-4* alleles (*ok837* and *lst1684*) at day 1 of adulthood. All data are normalized to the mean wild-type length. Small colored points represent individual worms, while large black circles denote the means of three independent biological replicates. Error bars represent ± SD. (D) Total progeny per individual hermaphrodite (wild-type vs. *cup-4*). Individual data points represent single hermaphrodites’ brood size. (E) Relative body length of wild-type and *cup-4* mutants at day 1 of adulthood. All data are normalized to the mean wild-type length. Small points represent individual worms, while large black circles denote the means of three independent biological replicates. Error bars represent ± SD.

**Fig. S3:**
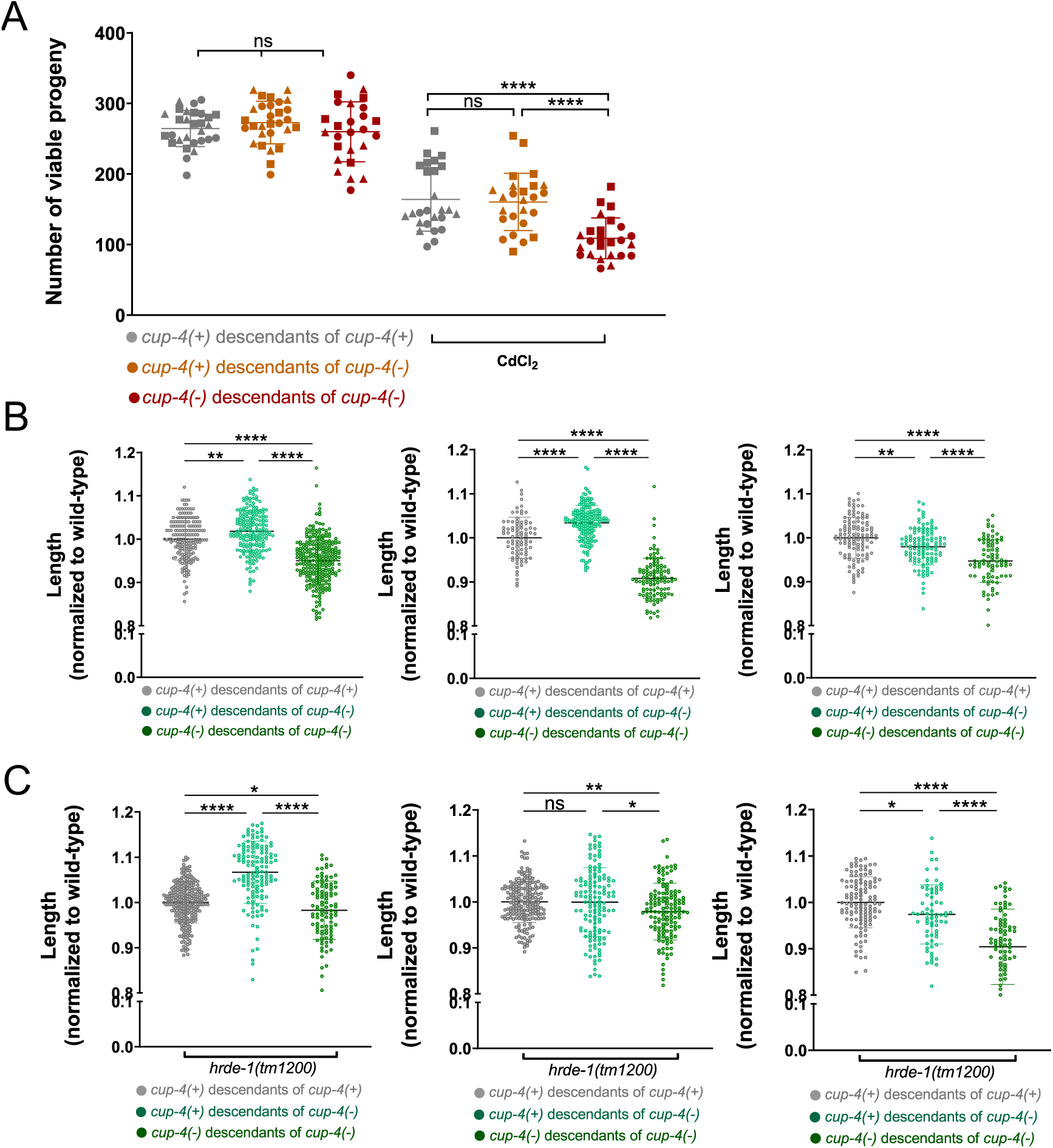
epigenetic inheritance of length and cadmium susceptibility. (A) Total progeny per individual cultured in the absence (left) or presence (right) of 50 µM CdCl₂. Individual data points represent single hermaphrodites’ brood size, with shapes indicating three independent experimental replicates. Data points for ‘*cup-4(+)* descendants of *cup-4(+)*’ and ‘*cup-4(-)* descendants of *cup-4(-)*’ are the same as shown in figure S2B. (B+C) Relative body length at day 1 of adulthood. All data are normalized to the mean ‘*cup-4(+)* descendants of *cup-4(+)*’ length. Shown are three biological replicates that show an inconsistent pattern. (C) worms containing a background mutation for *hrde-1*.

